# Single cell transcriptomics reveals that air-liquid interface culture promotes goblet cell differentiation and inhibits glycolysis in cell monolayers derived from rabbit caecum organoids

**DOI:** 10.1101/2025.07.21.665913

**Authors:** Tania Malonga, Emeline Lhuillier, Christelle Moreau, Elodie Riant, Cédric Cabau, Nathalie Vialaneix, Martin Beaumont

## Abstract

Faithfully recapitulating the cellular heterogeneity of the intestinal epithelium is essential when using organoid models. Air-liquid interface (ALI) culture has been shown to promote secretory cell differentiation but its impact on gene expression in each epithelial cell type remains unclear. In this study, we used single-cell RNA sequencing (scRNA-seq) to characterize the cellular heterogeneity of rabbit caecum-derived organoid monolayers grown under immerged or ALI conditions. We then compared these organoid cell type-specific gene expression profiles to a scRNA-seq atlas of the rabbit caecal epithelium *in vivo*. We selected the rabbit model notably because, unlike mice, it possesses BEST4+ epithelial cells, a newly discovered subset of mature absorptive cells. Our analysis revealed a high degree of transcriptomic similarity between *in vivo* and organoid-derived stem and transit-amplifying cells. ALI culture markedly enhanced the differentiation of the secretory lineage, especially goblet cells, which transcriptome closely resembled that of *in vivo* goblet cells. Furthermore, ALI was the only condition allowing the detection of enteroendocrine cells. BEST4+ cells, however, were absent from organoids in immerged or ALI conditions despite their presence *in vivo*. In addition, ALI culture led to a consistent downregulation of hypoxia and glycolysis-associated genes across all cell types, which suggests a metabolic shift likely driven by increased oxygen availability in ALI conditions. Cell-cell communication analyses further indicated that bone morphogenic protein (BMP) and fibroblast growth factor (FGF) signaling under ALI more closely mirrored *in vivo* patterns than under immerged condition. Altogether, these results demonstrate that ALI culture allows to better recapitulate the *in vivo* cellular heterogeneity and molecular signatures of the rabbit intestinal epithelium. Future optimization of culture conditions could enhance the physiological relevance of this organoid model, for instance by delivering oxygen exclusively to the basal side, as occurs *in vivo*.

## INTRODUCTION

The intestinal epithelium plays a crucial role in digestion and nutrient absorption, while protecting the organism from the external environment by acting as a physical and immunological barrier (1). These functions are performed by diverse epithelial cell types, all derived from stem cells located at the base of epithelial crypts (2). Epithelial progenitor cells migrate towards the crypt top and, after an initial phase of active proliferation (transit-amplifying [TA] cells), gradually differentiate towards the absorptive and secretory lineages (3). Single-cell RNA sequencing (scRNA-seq) recently identified subpopulations of absorptive cells, which includes the newly discovered BEST4^+^ cells specialized in ion secretion (4–7). Secretory cells include goblet cells secreting mucus, enteroendocrine cells producing hormones, Paneth cells secreting antimicrobial peptides, and Tuft cells involved in intestinal immunity (8, 9). Recapitulating this cellular diversity is critical when studying the intestinal epithelium *in vitro*.

Stem cell-derived intestinal organoids provide a three dimensional (3D) *in vitro* model able to recapitulate the cellular heterogeneity and differentiation dynamics of the intestinal epithelium (10). However, single cell transcriptomics showed that intestinal organoids cultured in expansion medium consist primarily of stem and progenitor cells, while mature differentiated cells remain rare or absent (11–13). Removal of mitogenic factors from the medium (e.g. Wnt ligands, epidermal growth factor) drive organoid cell differentiation, which can be oriented towards specific absorptive or secretory lineages by addition of cytokines or small molecules to the culture medium (7, 11, 12, 14–16). A major limitation of 3D intestinal organoids is the limited access to the apical side of epithelial cells, which is enclosed within the lumen. Alternatively, organoid cells can be dissociated to single cells and cultured as monolayers in inserts, which facilitates access to both apical and basal sides of epithelial cells (17). However, the cellular heterogeneity of organoid cell monolayers remains to be fully characterized, especially at the single cell transcriptomic level.

Removing the top medium of intestinal organoid cell monolayers to form an air-liquid interface (ALI) enhances differentiation, notably by increasing the population of goblet cells and mucus secretion (18–26). Furthermore, ALI culture also allows to recapitulate all stages of parasite infection in intestinal organoid epithelial cells (19, 27, 28). The mechanisms driving ALI-induced changes in epithelial phenotype are not fully understood, although probably involving metabolic adaptations upon higher oxygen supply (19, 29, 30).

In this study, we used scRNA-seq to understand how ALI culture affects the cellular heterogeneity of intestinal organoid cell monolayers. We compared rabbit caecum organoid cell monolayers cultured in submerged or ALI conditions and evaluated the similarity with an *in vivo* single cell transcriptome atlas of the rabbit caecum epithelium (31, 32). The rabbit was considered as a valuable model due to the presence of BEST4^+^ cells in the caecum *in vivo*, whereas these cells are lacking in mice (6). Furthermore, refinement of the culture condition of intestinal organoids from domestic animals is needed to improve the relevance of *in vitro* veterinary research, notably by recapitulating epithelial cell diversity (33).

## MATERIAL AND METHODS

### Isolation of epithelial crypts from the rabbit caecum

Epithelial crypts were isolated from the caecum of a 18-day-old suckling rabbit raised at the PECTOUL experimental facility (GenPhySE, INRAE, Toulouse, France). The protocol was approved by the local ethics committee (SSA n°115, SSA_2024_006V2). The caecum was isolated after euthanasia and placed in cold PBS (GIBCO, cat#10010-015). The caecum was then opened longitudinally and washed with cold PBS to remove all content. The tissue was minced into 0.5 cm^2^ sections before transfer to 5 mL of a pre-warmed (37°C) digestion solution prepared in HBSS without Ca^2+^/Mg^2+^ (ThermoFisher Scientific, cat#14175095) and supplemented with 5 mM EDTA (ThermoFisher Scientific, cat#AM9260G) and 1 mM DTT (Sigma, cat# 10197777001). After incubation (20 min at 37°C under slow agitation at 40 g), epithelial crypts were detached by vigorous manual shaking for one minute. The crypt solution was then filtered 100 µm to remove tissue fragments. The filtered crypt solution was centrifuged (300 g, 5 min, 4°C) and the pellet was resuspended in 5 mL cold PBS. Crypts were manually counted and the volume corresponding to approximately 900 crypts was centrifuged (300 g, 5 min, 4°C). The crypt pellet was resuspended in 1 mL freezing solution (80% DMEM [ThermoFischerScientific, cat#31966047], 10% fetal bovine serum [ThermoFischerScientific, cat#10270-106], 10% DMSO [Corning, cat#25-950-CQC], 10 µM Y27632 [StemCell Technologies, cat# 72304]) and transferred in a cryotube, which was placed in a CoolCell™ LX Cell Freezing Container (Corning, cat# 432003) at −80°C for 24 h before long-term storage in liquid nitrogen.

### Culture of rabbit caecum organoids in three dimensions

Cryopreserved rabbit caecum epithelial crypts were thawed at 37°C before centrifugation (300 g, 5 min, room temperature). The crypt pellet was resuspended in Matrigel (Corning, cat#354234), plated into a pre-warmed 48-well plate (37°C) (25 µL/well), and left to polymerize for 15 min at 37°C. Each well was overlaid with 250 µL of organoid growth medium composed of IntestiCult Organoid Growth Medium (Human) (Stem Cell Technologies, cat# 06010) supplemented with 1% penicillin/streptomycin (PS, Sigma, cat#P4333), and 100 µg/mL Primocin (InvivoGen, cat#ant-pm-05), at room temperature. The plate was placed in a cell culture incubator (37°C, 5% CO_2_) and the medium was changed every 2-3 days.

Organoids were passaged 7 days after crypt seeding. After a wash in warm PBS, organoids in Matrigel domes were homogenized in pre-warmed TrypLE (Gibco, cat# 12605-010). After incubation for 5 min at 37°C in a CO_2_ incubator, the Matrigel-cell suspension was homogenized by pipetting and the incubation step at 37°C was repeated once. Digestion was stopped by adding DMEM supplemented with 10% FBS and 1% PS (DMEMc). The contents of each well were pooled into a tube and centrifuged (500 g, 4°C, 5 min). Cells were resuspended in DMEMc and counted with a using a Countess 3 Automated Cell Counter (ThermoFischerScientific, cat#16842556). Organoid cells were resuspended in cold Matrigel:DMEMc (v/v: 2:1), before plating into pre-heated (37°C) 24-well plates (3000 cells/50 µL/well). The plates were left to polymerize for 30 min at 37°C. Each well was then overlaid with 500 µL of organoid growth medium at room temperature. The plate was placed in a cell culture incubator (37°C, 5% CO_2_) and the medium was changed every 2-3 days.

### Culture of cell monolayers derived from the rabbit caecum organoids

Cell culture inserts for 24-well plates (Corning, cat#353095) were coated with 150 µL of 50 µg/mL type IV collagen derived from human placenta (Sigma, cat# C5533-5MG) at 37°C for 2 hours. After removal of the coating solutions, inserts were dried without the lid under the cell culture cabinet until seeding. Nine days after passaging in 3D, the Matrigel domes with organoids were dissociated by pipetting and transferred into tubes containing 5 mL of cold DMEMc. The suspension was centrifuged (500 g, 4°C, 5 min), and the supernatant above the Matrigel layer was disrupted by pipetting. Then, organoids were centrifuged (500 g, 4°C, 5 min) and the resulting pellet was resuspended in pre-warmed TrypLE supplemented with 10 µM Y27632 and incubated in a 37°C water bath for 5 min before homogenization by pipetting. This cycle of incubation and homogenization was repeated until complete cell dissociation. Cold DMEMc was added to the suspension to stop digestion before centrifugation (500 g, 4°C, 5 min).

The cell pellet was resuspended in DMEMc and cells were counted as described before. Cells were resuspended in organoid growth medium supplemented with 20% FBS and 10 µM Y27632 before seeding in collagen IV-coated inserts (2.5 10^5^ cells/ 200 µL/insert). The same culture medium was also added to the basal side (500 µL). Cells were incubated at 37°C under 5% CO_2_ atmosphere. Three days after seeding, the culture medium was removed from the basal and apical sides and cell monolayers were washed with PBS. The apical compartment was either submerged in 200 µL warm DMEM supplemented with PS 1% on the apical side (control condition) or remained empty to set up an air-liquid interface (ALI condition). Organoid culture medium supplemented with 20% FBS was added to the basal side (500 µL) in both control and ALI conditions. Four days after seeding, apical (control condition) and basal (control and ALI conditions) media were refreshed. Five days after seeding (*i.e.,* after 48h of culture in control or ALI conditions), the cell monolayers were processed for single cell transcriptomics.

### Organoid cell monolayer dissociation and multiplexing

Cells dissociation was performed on n = 4 cell culture inserts per condition (control or ALI). Cell monolayers were washed in PBS before adding pre-warmed TrypLE supplemented with 10 µM Y27632 in the apical (200 µL) and basolateral (500 µL) compartments. Cells were incubated at 37°C for 20 min with homogenization every 5 min by pipetting. The dissociated cells were transferred into 1.5 mL tubes containing 1 mL of cold DMEMc to stop digestion and centrifuged (500 g, 5 min, 4°C). The pellet was resuspended in 100 µL of cold PBS containing 10% FBS and 10 µM Y27632 before counting. Approximately 10^5^ cells/insert were resuspended in 1 mL with cold PBS containing 10% FBS and 10 µM Y27632 before centrifugation (300 g, 5 min, 4°C). The supernatant was removed and 100 µL of a single Cell Multiplexing Oligo (CMO) solution from the 3’ CellPlex Kit Set A (10X Genomics, cat#1000261) was addeed to each tube (1 CMO/insert). The cells were homogenized with the CMO solution by pipetting 15 times before incubation at room temperature for 5 min. Subsequently, 1.4 mL of cold PBS containing 10% FBS and 10 µM Y27632 was added before mixing by pipetting and centrifugation (300 g, 5 min, 4°C). The supernatant was discarded and cells were resuspended in 100 µL of cold PBS containing 10% FBS and 10 µM Y27632 before pooling cells derived from inserts treated in the same condition (control or ALI) in equal proportions (1:1:1:1, n = 4 inserts pooled per condition).

### Cell sorting and library construction

For viability assessment, cells were centrifuged (300 g, 5 min, 4°C) and resuspended in 1 mL of PBS supplemented with 10 µM Y27632 and 1 µL of LIVE/DEAD™ Fixable Violet Dead Cell Stain (ThermoFisher Scientific, cat#L34963). After incubation (4°C, 30 min, in the dark), cells were centrifuged (300 g, 5 min, 4°C) and the pellet was washed twice with FACS buffer (3% FBS, 2 mM EDTA, 10 µM Y27632 in PBS) before filtering (40 µm). Approximately 10^5^ live cells were sorted by using a BD Influx cell sorter instrument with a 100 µm nozzle, under 20 psi at the I2MC Cytometry and Cell sorting TRI platform (Toulouse, France). Cells were centrifuged (300 g, 10 min, 4°C), resuspended in PBS and manually counted.

For each sample (control or ALI), 50,000 cells were used for encapsulation into droplets using Chromium NextGEM Single Cell 3′ Kit v3.1 (10X Genomics, cat#PN-1000268) with Feature Barcoding technology for Cell Multiplexing, according to manufacturer’s protocol (10x Genomics CG000388 Rev C user guide). Briefly, after generation of Gel bead-in-EMulsions (GEMs) using Chromium Next GEM Chip G, GEMs were reverse transcribed in a C1000 Touch Thermal Cycler (BioRad) programmed at 53°C for 45 min, 85°C for 5 min, and held at 4°C to produce barcoded cDNA from polyA mRNA and barcoded DNA from the CMO. Then, single-cell droplets were broken and cDNA was isolated and cleaned with Cleanup Mix containing DynaBeads (ThermoFisher Scientific). cDNA was then amplified by PCR with a C1000 Touch Thermal Cycler programmed at 98°C for 3 min, 11 cycles of (98°C for 15 s, 63°C for 20 s, 72°C for 1 min), 72°C for 1 min, and held at 4°C. cDNA derived from mRNA or CMO were separated thanks to differential clean up with SPRIselect beads (Beckman Coulter, cat# B23317). 3’ Gene Expression Library was constructed with approximately 80-90 ng of amplified cDNA, which was fragmented, end-repaired, A-tailed, index adaptor ligated, and cleaned with SPRIselect beads in between steps. Post-ligation product was amplified and indexed with a C1000 Touch Thermal Cycler programmed at 98°C for 45 s, 13 cycles of (98°C for 20 s, 54°C for 30 s, 72°C for 20 s), 72°C for 1 min, and held at 4°C. Cell Multiplexing Libraries were constructed and indexed by PCR too, using similar parameters : 98°C for 45 s, 6 cycles of (98°C for 20 s, 54°C for 30 s, 72°C for 20 s), 72°C for 1 min, and held at 4°C. The sequencing-ready libraries were cleaned up with SPRIselect beads. Libraries were pooled following the recommendations, and loaded with 1% PhiX on two S1 lanes of the NovaSeq 6000 instrument (Illumina) using the NovaSeq 6000 S1 Reagent Kit v1.5 (100 cycles), and the following sequencing parameters: 28 bp read 1 – 10 bp index 1 (i7) – 10 bp index 1 (i5) – 88 bp read 2.

### scRNA-seq pre-processing, filtering, normalization and clustering

Cell Ranger software (version 7.1.0, 10x Genomics) was used to align and quantify the raw sequencing data using the rabbit genome (GCF_009806435.1_UM_NZW_1.0). A custom reference file was generated using the mkgtf command with the parameters ‘--attribute=gene_biotype:protein_coding’ and ‘-- attribute=gene_biotype:lncRNA’, followed by the mkref command with default parameters. The multiplexed data was then analyzed using the multi pipeline, run with default parameters.

Using R software (version 4.2.1), the Seurat pipeline (version 4.3.0; Butler et al. 2018) was employed for preprocessing and analysis. Control and ALI datasets were preprocessed independently. The count matrices of each sample (cells derived from a single culture insert) of a single condition (control or ALI) were merged. Cells expressing fewer than 1,500 genes, more than 8,000 genes, or exceeding 50,000 RNA counts were filtered out. Additionally, cells with mitochondrial reads content exceeding 20% of total reads were removed. The data were then normalized using the Seurat *NormalizeData* function with the *LogNormalize* method. The top 3,000 highly variable genes were identified using the *FindVariableFeatures* function, based on a mean-variance trend. Principal Component Analysis (PCA) was performed using these variable genes, retaining 30 Principal Components (PC) for downstream analyses. The Seurat *CellCycleScoring* function was used to score and assign phases of the cell cycle to each cell. PC were used as features to cluster cells using the Leiden algorithm applied to a nearest-neighbor graph, with a resolution of 0.9 for the control dataset and 0.6 for the ALI dataset: The resolution was chosen independently in each condition as the minimal value to allow obtaining separate clusters for stem cells (S cycle phase) and TA cells (G2M cycle phase). The Uniform Manifold Approximation and Projection (UMAP) method was also run on PC for dimensionality reduction and visualization.

### Cell type assignment

#### Marker genes

To identify cluster-specific markers, we then used the *FindMarkers* function. For every cluster, all genes expressed in a minimum of 25% of the cells of this cluster were subjected to a Wilcoxon test for differential expression between this cluster and all the other clusters. *P*-values were corrected for multiple testing using Bonferroni correction. Genes were selected as markers if their adjusted *p*-value was below 0.05 and if they were over-expressed in the cluster of interest compared to the others. Further filtering was performed to ensure that only genes with a log-Fold Change (logFC) of at least 0.25 between the cluster of interest and the other clusters were considered. Cell types were then manually assigned to each cluster based on the identified markers and existing cell type references from rabbit, pig and human intestinal epithelium (8, 32, 35–37).

#### Automatic assignation of cell types

To validate the manual cell type annotation, an automatic labeling approach was employed by transferring annotations from a reference *in vivo* dataset of rabbit caecum epithelial cells (32, 38). The same preprocessing steps as for the organoid dataset were performed on the *in vivo* reference dataset. Then, the *FindTransferAnchors* function was used on the first 30 PC to identify cell pairs, or “anchors,” between the organoid and *in vivo* datasets. These anchors were then used with the *MapQuery* function to map the organoid cells into the *in vivo* rabbit caecum epithelial cell space. The reference annotations were subsequently transferred to the organoid dataset and visualized using the UMAP embedding. Additionally, the results of *MapQuery* were used to compute a mapping score (function *MappingScore*) for each organoid cell, indicating how strongly each cell neighborhood is aligned with the reference dataset (a higher score represents a closer match to the reference).

#### Crypt axis gene score

The crypt axis gene (CAG) score of each cell was calculated via the *AddModulesScore* function by averaging the expression of genes previously defined as expressed in epithelial cells located at the crypt top (*PLAC8, CEACAM1, TSPAN1, DHRS9, PKIB, HPGD*) (8).

#### Biological pathway enrichment

First, the marker genes of each cell type were identified as described in the *Marker genes* section. Then, marker genes for each cell type were subjected to biological enrichment analysis using the *enrichGO* function from the *clusterProfiler* package (version 4.6.1; Yu et al. 2012), with the entire set of expressed genes used as the reference background. As there is no *Oryctolagus cuniculus* (rabbit) database, *Homo sapiens* was used as the reference species for the enrichment analysis. Redundancy in the results was reduced by the application of the *simplify* function from *clusterProfiler*, which removed terms with a semantic similarity greater than 0.7 and retained the most significant terms (with the smallest *p*-value) in each term group. Multiple testing correction was performed using the Benjamini-Hochberg (BH) method (40), and pathways were considered significantly enriched if the adjusted *p*-value was less than 0.05.

#### Data integration of *in vitro* datasets and visualization

To compare the two culture conditions (control and ALI), both datasets were integrated into a single dataset using the *IntegrateData* function. Integration features were selected using the *SelectIntegrationFeatures* function, and integration anchors were identified using the *FindIntegrationAnchors* function. Integrated data were scaled with *ScaleData*. PCA was conducted on scaled integrated data and cells were then visualized on the UMAP embedding obtained from the first 30 PC.

### Differential analysis of gene expression

Each cell type was analyzed independently. Genes that were expressed in less than 30% of the cells were filtered for the differential analysis only. Pseudo-counts for each gene were then calculated by summing the raw gene counts from cells of the same sample for each cell type. This step is considered crucial, as it has been demonstrated that differential analysis on pseudo-bulk data produces more robust results, minimizing Type I errors compared to direct analysis of scRNA-seq data (41, 42). The pseudo-counts were normalized across samples for each cell type using edgeR’s “TMM” method (43). Differential expression analysis was performed using a Negative Binomial generalized linear model in *edgeR*, with gene expression explained by a fixed effect of the culture condition. *P*-values were calculated using a log-likelihood ratio (LR) test for the culture condition effect, and adjusted *p*-values were derived via the BH method. Genes with adjusted *p*-values < 0.05 were considered differentially expressed in the cell type of interest. These genes were further analyzed for enrichment in biological pathways, as described in the “Biological pathway enrichment” section.

### Comparison of the *in vivo* and *in vitro* datasets

To compare the organoid (control and ALI) and *in vivo* datasets, the three datasets were integrated into a single dataset using the *IntegrateData* function. Integration features were selected using the *SelectIntegrationFeatures* function, and integration anchors were identified using the *FindIntegrationAnchors* function. Integrated data were scaled with *ScaleData*. PCA was conducted on scaled integrated data and cells were then visualized on the UMAP embedding obtained from the first 30 PC.

The miloR (44) package was used to compare the similarity of cell neighborhoods by cell type between organoid datasets (control and ALI conditions) and *in vivo* data. Each Seurat object was first converted into a *SingleCellExperiment* object and then transformed into a *Milo* object to enable neighborhood analysis. The construction of neighborhoods was carried out in multiple steps. First, within the PCA space, the *k* = 10 nearest neighbors for each cell were obtained and a *k*-nearest neighbor graph was derived. Vertices (*i.e.*, cells) and corresponding neighborhoods were then randomly selected (*prop = 0.1*) with the *makeNhoods* function using a refined strategy to improve selection and neighborhood representation (*refined = TRUE*). The *calcNhoodSim* function was used to compute the Pearson correlation between neighborhoods across the two conditions, using a maximum of 2,000 highly variable genes in the analysis. The distribution of neighborhood correlations between organoid cells (control and ALI) and the *in vivo* datasets is visualized using *plotNhoodSimGroups* for each cell type of the organoid.

### Cell-cell communication

Cell-cell communication analysis was performed using the CellChat R package (version 2.1.2; Jin et al. 2021; Jin, Plikus, et Nie 2023). All the functions of CellChat were used with their default parameters. To identify interactions between cell populations, we first identified significantly over-expressed ligands and receptors in each cell group using the Wilcoxon test implemented in the *identifyOverExpressedGenes* and *identifyOverExpressedInteractions* functions with default options. This allowed to identify ligand-receptor pairs where either ligand or receptor was significantly expressed. Biologically relevant interactions were estimated using the *computeCommunProb* function, which estimates communication probabilities by integrating gene expression data with known ligand-receptor interactions. Using the *computeCommunProbPathway* function, CellChat calculates the cell-cell communication probability at the signaling pathway level by aggregating the communication probabilities of all ligand-receptor interactions related to each signaling pathway. By default, CellChat employed the *type = “13rimean”* method to calculate the average gene expression per cell group, prioritizing strong interactions while minimizing false positives. To account for differences in population sizes across cell groups, the probability calculation incorporated cell proportions by setting *population.size = TRUE*. Interactions were then filtered to retain those involving at least ten cells per group (*min.cells = 10*) using the *filterCommunication* function. The identified ligand-receptor interactions were extracted as a data frame using *subsetCommunication*, enabling visualization at the level of individual interactions or at the level of the entire signaling pathways.

### Data availability

The scRNA-seq organoid data for this study have been deposited in the European Nucleotide Archive (ENA) at EMBL-EBI under accession number PRJEB88083 (www.ebi.ac.uk/ena/browser/view/PRJEB88083). The organoid data are also accessible on the FAANG portal (https://data.faang.org/dataset/PRJEB88083). The *in vivo* scRNA-seq data of the rabbit caecum epithelium are accessible at EMBL-EBI under accession number PRJEB74645 (www.ebi.ac.uk/ena/browser/view/PRJEB74645) and on the FAANG portal (https://data.faang.org/dataset/PRJEB74645) and on the searchable Broad Institute Single-cell Portal (https://singlecell.broadinstitute.org/single_cell/study/SCP2662/single-cell-transcriptomics-in-caecum-epithelial-cells-of-suckling-rabbits-with-or-without-access-to-solid-food).

## Results

The aim of this study was to assess the impact of ALI culture on the single cell transcriptome of cell monolayers derived from rabbit caecum organoids (Figure 1A). Half of the monolayers were cultivated in submerged conditions (control group, n = 4 culture inserts) and the other half were cultivated in ALI condition (ALI group, n = 4 culture inserts).

**Figure 1:**
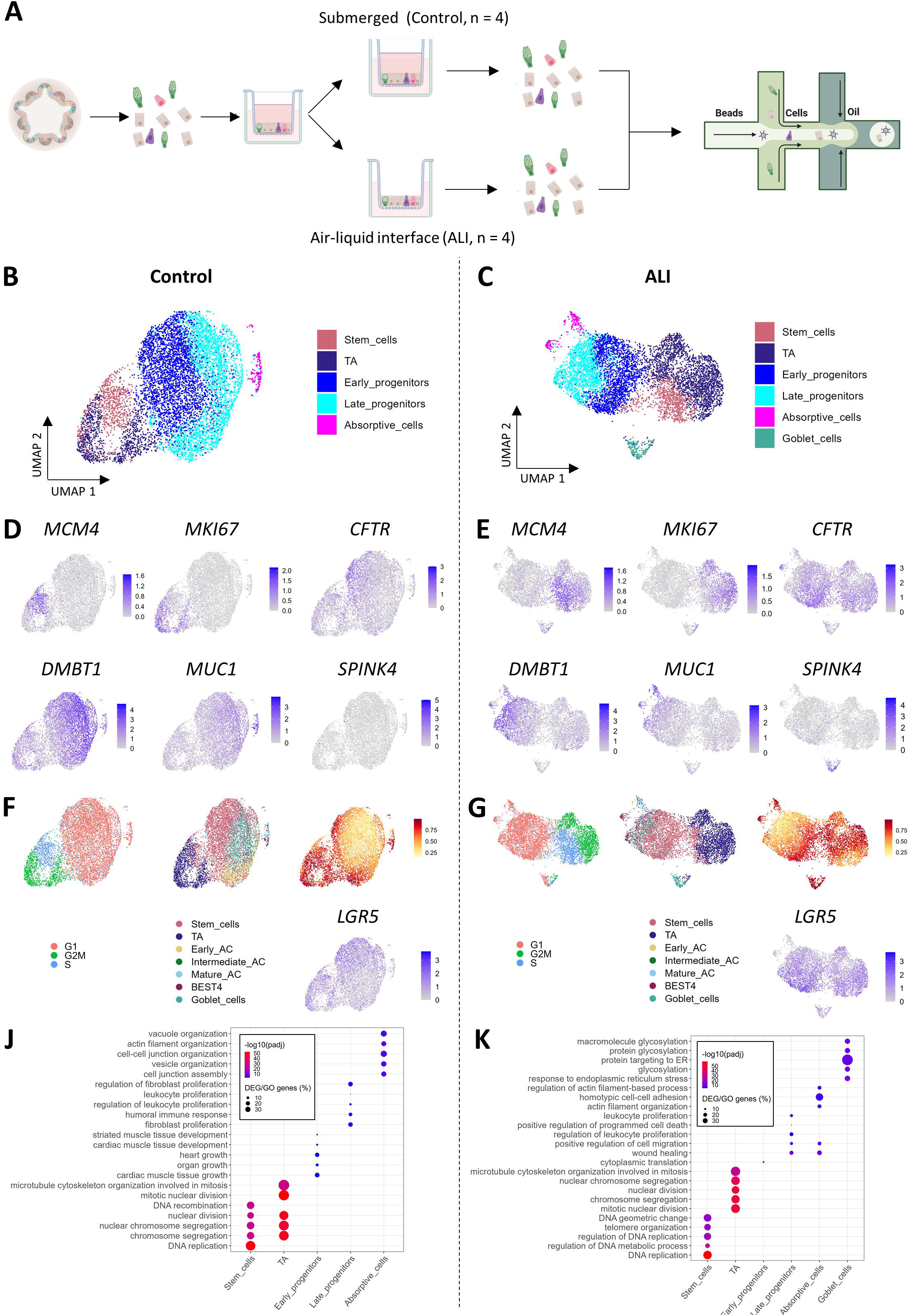
A single-cell transcriptomic atlas of intestinal organoid cell monolayers cultivated in submerged or air-liquid interface conditions. (A) Experimental workflow. Rabbit 3D caecum organoids were dissociated to single cells and seeded in inserts to form cell monolayers. Air-liquid interface (ALI) was created by removing the apical medium from the top chamber for 2 days. Cell monolayers cultured in submerged (control, n = 4 inserts) or ALI (n = 4 inserts) were dissociated and analyzed by droplet-based single cell transcriptomics. (B) and (C) Uniform Manifold Approximation and Projection (UMAP) of cells colored by epithelial cell types in the submerged (control) and ALI conditions, respectively. (D) and (E) UMAPs colored by the expression of the stem cell marker *MCM4*, the TA cell marker *MKI67*, the early progenitor cell marker *CFTR*, the late progenitor cell marker *DMBT1*, the absorptive cell marker *MUC1* or the goblet cell marker *SPINK4* in the immerged (left panel) and ALI conditions cells (right panel), respectively. (F) and (G) UMAP colored by the inferred cell cycle state (left panel), the cell types assigned through automatic annotation based on an *in vivo* rabbit caecum epithelial cell atlas (middle panel), the mapping score (right panel), and the expression of the *LGR5* stem cell marker gene (downright panel) for cells from the control and ALI conditions, respectively. (J) and (K) Top five biological processes (ranked by adjusted *p*-value) enriched in marker genes for each cell type from the control and the ALI conditions, respectively. Color represents the -log10(adjusted *p*-value) from the over-representation test, while size indicates the percentage of marker genes among the genes associated with this ontology term. AC: absorptive cells, TA: transit-amplifying cells.

### Cellular diversity in monolayers derived from caecum organoids

After filtering the cells using quality parameters, the control dataset and the ALI dataset included 7,888 and 7,138 cells, respectively (Figure S1A). Each sample contained an equivalent number of cells in each condition (Figure S1B), and the quality parameters were similar across experimental conditions (Figure S1C). We identified 11 cell clusters in the control condition and 12 cell clusters in the ALI condition (Figure S1D). Assignation of cell types to each cluster was performed using known markers of intestinal epithelial lineages (Figures 1B-E), cell cycle score prediction, and automatic annotation using transfer of cell labels from *in vivo* rabbit caecum epithelium (Figures 1F and G). Assignation of cell types was performed separately for each condition to allow the identification of condition-specific cell populations that might otherwise be overlooked in an integrated analysis. Biological processes enriched in each cell type were predicted based on their marker genes (Figures 1J and K). The lists of marker genes and of enriched biological processes are presented for each cell type in each condition in Tables S1-S2 and Tables S3-S4, respectively.

In both control and ALI conditions, stem cells were identified as *MCM*4^+^ and *MCM5*^+^ and were predicted in the S phase of the cell cycle (Figures 1B-G). As expected, stem cell marker genes were mostly involved in DNA replication (Figures 1J and K). Transit amplifying (TA) cells were identified as *MKI67*^+^, *TOP2A*^+^, and *UBE2C*^+^, and were predicted in the G2M phase of the cell cycle (Figures 1B-G). Marker genes of TA cells were enriched in genes involved in mitosis (Figures 1J and K), reflecting their active proliferation. Almost all the other cells were predicted to be in the G1 phase of the cell cycle (Figure 1F and G). Early progenitors highly expressed *CFTR* (Figures 1B-E). Late progenitors were identified by *DMBT1* expression (Figures 1B-E) and their marker genes were involved in immune response (Figures 1J and K). Absorptive cells were identified as *MUC1*^+^ and *SELENOP*^+^ (Figures 1B-E). Functions enriched in marker genes of absorptive cells included “actin filament organization,” “vacuole organization,” and “cell-cell junction organization” (Figures 1J and K), which are crucial for the transport and barrier functions of these cells. Goblet cells, which were only present in the ALI dataset, were identified as *ATOH1*^+^, *SPINK4*^+^, *FCBGP*^+^, and *TFF3*^+^ (Figure 1B-E). The presence of S and G2M cells in the goblet cell cluster suggested the presence of dividing goblet cell progenitors in the ALI organoid cell monolayers (Figure 1G). Marker genes of goblet cells were involved in “glycosylation” and “response to endoplasmic reticulum stress” (Figure 1K), which is consistent with the role of goblet cells in mucus secretion.

The manual identification of stem cells, TA cells, mature absorptive cells, and goblet cells was confirmed by the automatic assignation, which was associated with the high mapping score for these cell types (Figure 1F and G). The presence of BEST4^+^ cells was predicted by automatic annotation in both control and ALI conditions but this labeling was not confirmed by manual annotation as these cells did not express the canonical markers *BEST4*, *OTOP2*, or *CA7*. Automatic annotation also identified a high proportion of stem cells in the clusters manually identified as early and late progenitors, although these cells were predicted in the G1 phase and had a low mapping score. This overestimation of stem cell proportion by the automatic annotation may be driven by the widespread expression of the stem cell marker *LGR5* in organoid cells (Figure 1F and G).

### Comparison of the transcriptomic profiles of each cell type between the control and ALI conditions

Globally, the marker genes of each cell type were highly conserved across the two conditions (Figures 2A and B). Accordingly, integration of control and ALI datasets revealed that cells derived from each condition co-localized by cell type in the UMAP of the integrated data (Figure 2C and D), which suggests an overall transcriptomic similarity between cells derived from each culture condition. In contrast, goblet cells were derived exclusively from the ALI dataset in which they composed about 3% of the cells (Figure 2C-E). The proportion of stem and TA cells was higher in the ALI condition when compared to the control condition, while the opposite was observed for progenitor cells (Figure 2E). The percentage of absorptive cells was similar in the two conditions and represented less than 3% of cells (Figure 2E). The crypt axis gene score in each cell type was consistent with their expected positions and was equivalent in the two conditions (Figure 2F).

**Figure 2:**
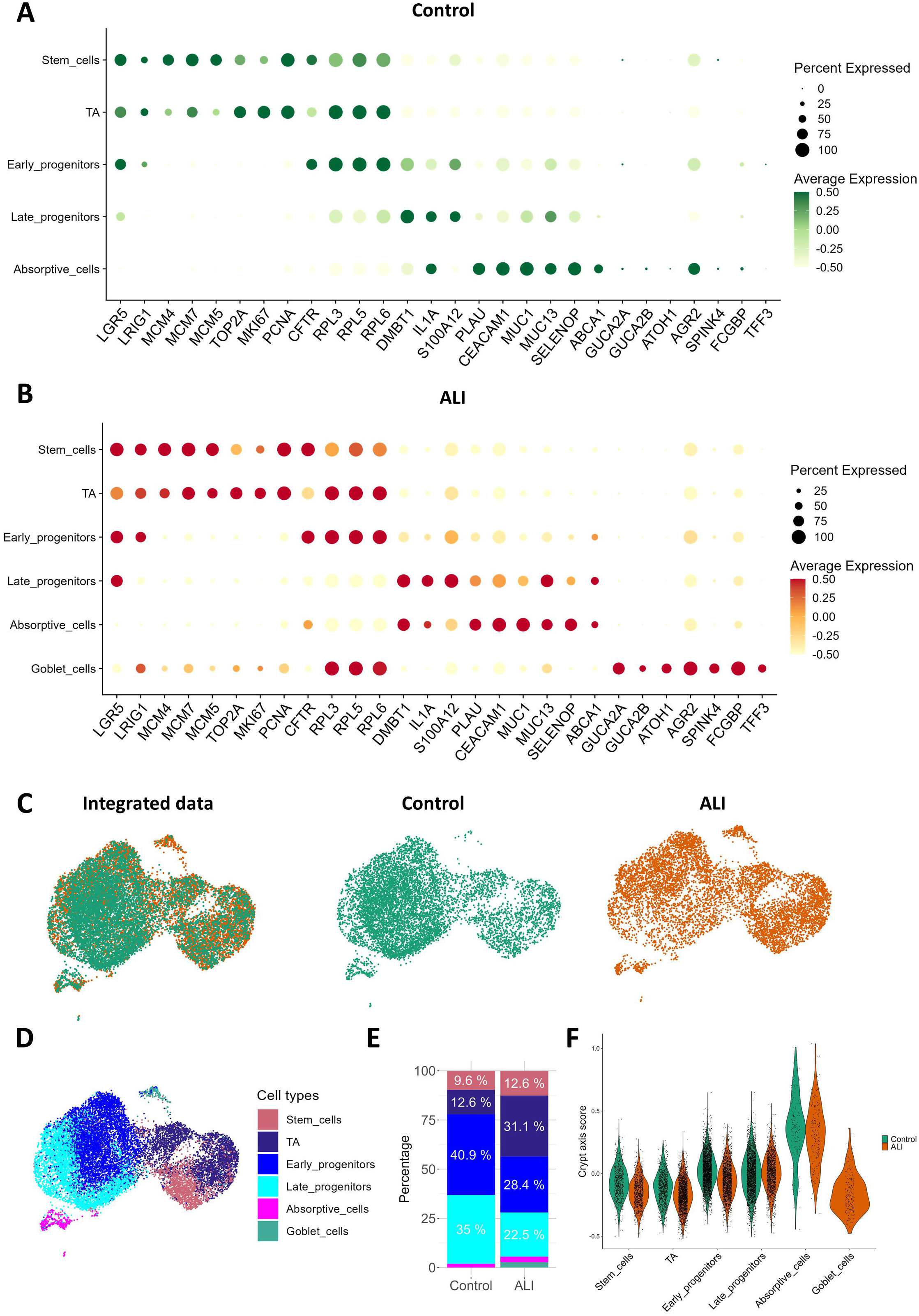
Comparison of the single cell transcriptome of organoid cell monolayers cultured in submerged or air-liquid interface conditions. (A) and (B) Expression of selected marker genes for each cell type from the submerged (control) and the air-liquid interface (ALI) conditions. The size corresponds to the percentage of cells expressing the gene in the cell type. The color intensity corresponds to the average scaled expression. (C) Uniform Manifold Approximation and Projection (UMAP) of integrated datasets colored by culture condition. The left panel displays a UMAP of both datasets, with cells cultivated in control (n = 4) and ALI (n = 4) conditions. Cells from the control and ALI conditions are shown independently in the middle and right panels, respectively. (D) UMAP of integrated data colored by epithelial cell types. (E) Relative abundance of each cell type per condition. (F) Crypt axis score for each cell type per condition. TA: transit-amplifying cells.

A differential analysis was performed to compare the expression of genes between the two conditions in each cell type, except for goblet cells, which were present only in the ALI condition. Furthermore, a biological enrichment analysis was performed using the differentially expressed genes (DEG) to determine biological functions modulated by the culture condition in each cell type. The lists of DEGs and biological processes enrichment analysis are presented for each cell type in Tables S5 and S6, respectively. “Cellular response to hypoxia” was enriched in DEGs identified in absorptive cells (Figure 3A). Indeed, genes involved in hypoxia (*EGLN3*, *BNIP3, ERO1A, HMOX1*) were strongly downregulated by ALI culture in absorptive cells and in all other cell types (Figure 3B). “Glycolytic process” was enriched in absorptive cell DEGs (Figure 3A). Accordingly, the ALI culture strongly downregulated several genes coding for enzymes involved in glycolysis, including *GPI, ALDOA, TPI1, GADPH, PGK1, PGAM1*, and *ENO1,* in absorptive cells and in all other cell types (Figure 3C). Furthermore, ALI culture regulated the expression of genes involved in “ATP metabolic process” and “ATP generation from ADP” in late progenitors and absorptive cells, respectively (Figure 3A). Overall, these results indicate that ALI culture remodeled energy metabolism in cell monolayers derived from organoids.

**Figure 3:**
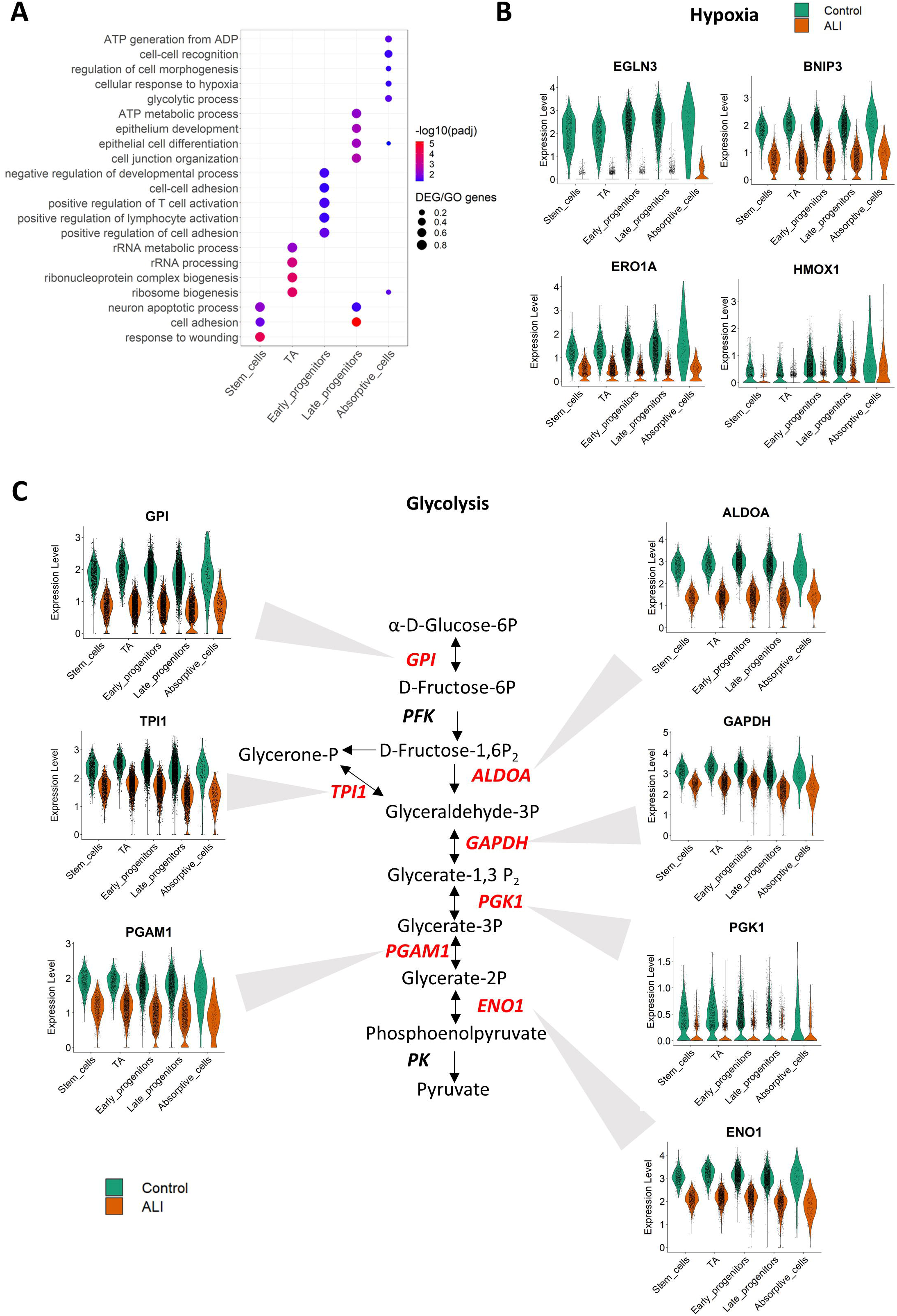
Single cell transcriptomics changes induced by culture in air-liquid interface across all cell types in organoid cell monolayers. (A) Selected biological processes enriched in differentially expressed genes (DEGs) per cell type according to the culture in submerged (control) or air-liquid interface (ALI) conditions. The color corresponds to the -log10(adjusted *p*-value) and the size represents the percentage of DEGs included in the biological process. (B) Expression level of DEGs involved in the hypoxia response, for each cell (dot) per cell type and condition. (C) Expression levels of selected DEGs involved in glycolysis across all cell types and conditions, with each cell represented as dots per cell type. A simplified representation of glycolysis enzymatic reactions is shown with enzyme-coding DEGs highlighted in red. TA: transit-amplifying cells.

### Comparison of the transcriptomic profiles of each cell types in organoids with *in vivo* epithelial cells

Organoid cell monolayer scRNA-seq data from the two culture conditions (control and ALI) were integrated with an *in vivo* rabbit caecum epithelium scRNA-seq dataset to compare the transcriptomic profiles of each cell type. Organoid stem cells, TA, and goblet cells mostly co-localized with their *in vivo* counterparts in the UMAP of integrated data (Figure 4A). Accordingly, the expression patterns of markers of these cell types (*MCM4* for stem cells, *TOP2A* for TA cells, and *SPINK4* for goblet cells) was similar across *in vivo* and organoid datasets (Figure 4B-D). In contrast, early progenitors, late progenitors and absorptive cells derived from organoids did not co-localize with their *in vivo* counterparts in the UMAP, except for rare cells (Figure 4A). Despite this overall transcriptomic divergence in the absorptive lineage suggested by UMAP representation, organoid progenitor and absorptive cells shared marker genes with absorptive *in vivo* cells, like *CFTR* for early progenitors, *DMBT1* for late progenitors and *SELENOP* for absorptive cells (Figure 4E-G). A few cells derived from organoid cell monolayers culture in ALI co-localized in the UMAP with enteroendocrine cells (EEC) and expressed canonical markers of this cell type (e.g. *CHGA, CHGB*) (Figure S2). In contrast, no organoid cells co-localized with the *in vivo* BEST4^+^ cells (Figure 4A).

**Figure 4:**
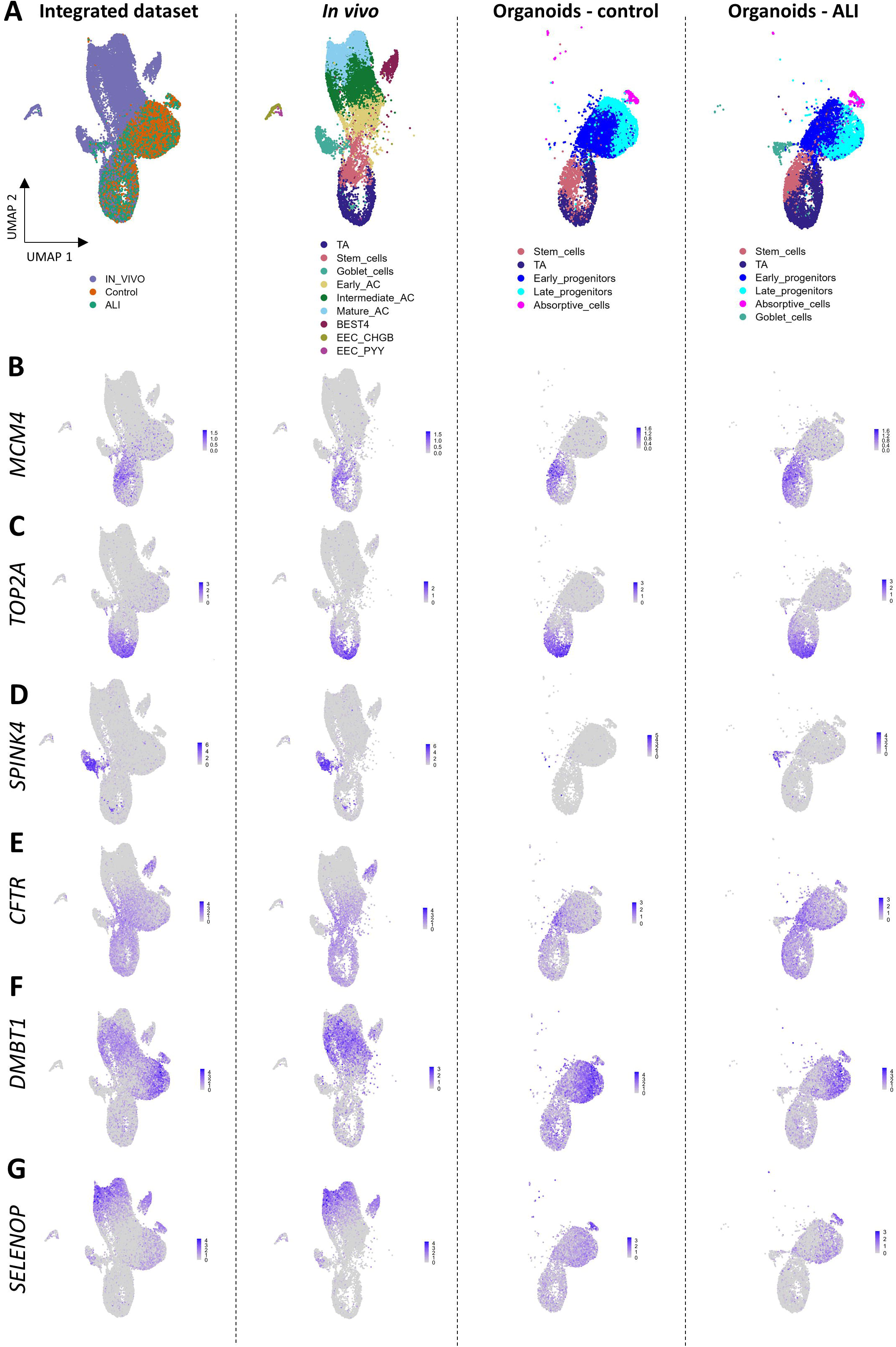
Comparison of the single cell transcriptome of caecum epithelial cells *in vivo* and in organoid cell monolayers. (A) Uniform Manifold Approximation and Projection (UMAP) of integrated single cell transcriptomics datasets of caecum epithelial cells *in vivo* and in organoid cell monolayers cultured submerged (control) or under air-liquid interface (ALI), colored by condition. The left panel displays the UMAP of integrated datasets, which include cells from all conditions. The other panels display the UMAP of integrated datasets with cells restricted to each group (*in vivo*, control organoid cells, ALI organoid cells) and colored by cell type. (B – G) UMAP of integrated datasets colored by the expression of (B) the stem cell marker *MCM4*, (C) the TA cell marker *TOP2A*, (D) the goblet cell marker *SPINK4*, (E) the progenitor cell marker *CFTR*, (F) the intermediate absorptive cell marker *DMBT1*, and (G) the mature absorptive cell marker *SELENOP* with all cells (left panel) or restricted to cells from each condition (other panels). AC: absorptive cells, EEC: enteroendocrine cells, TA: transit-amplifying cells.

Cell neighborhood correlation analysis revealed that the transcriptome of goblet cells from the ALI condition had the highest similarity with *in vivo* data (correlations ranging 0.75 to 0.9 across cells), followed by stem cells (0.65 to 0.9), and TA cells (0.7 to 0.85) (Figure 5A). The broad distribution of the crypt axis gene score from the *in vivo* cells reflected their distribution in all the crypt length, while the organoid cells were predicted to be positioned mostly in the crypt base (Figure 5B).

**Figure 5:**
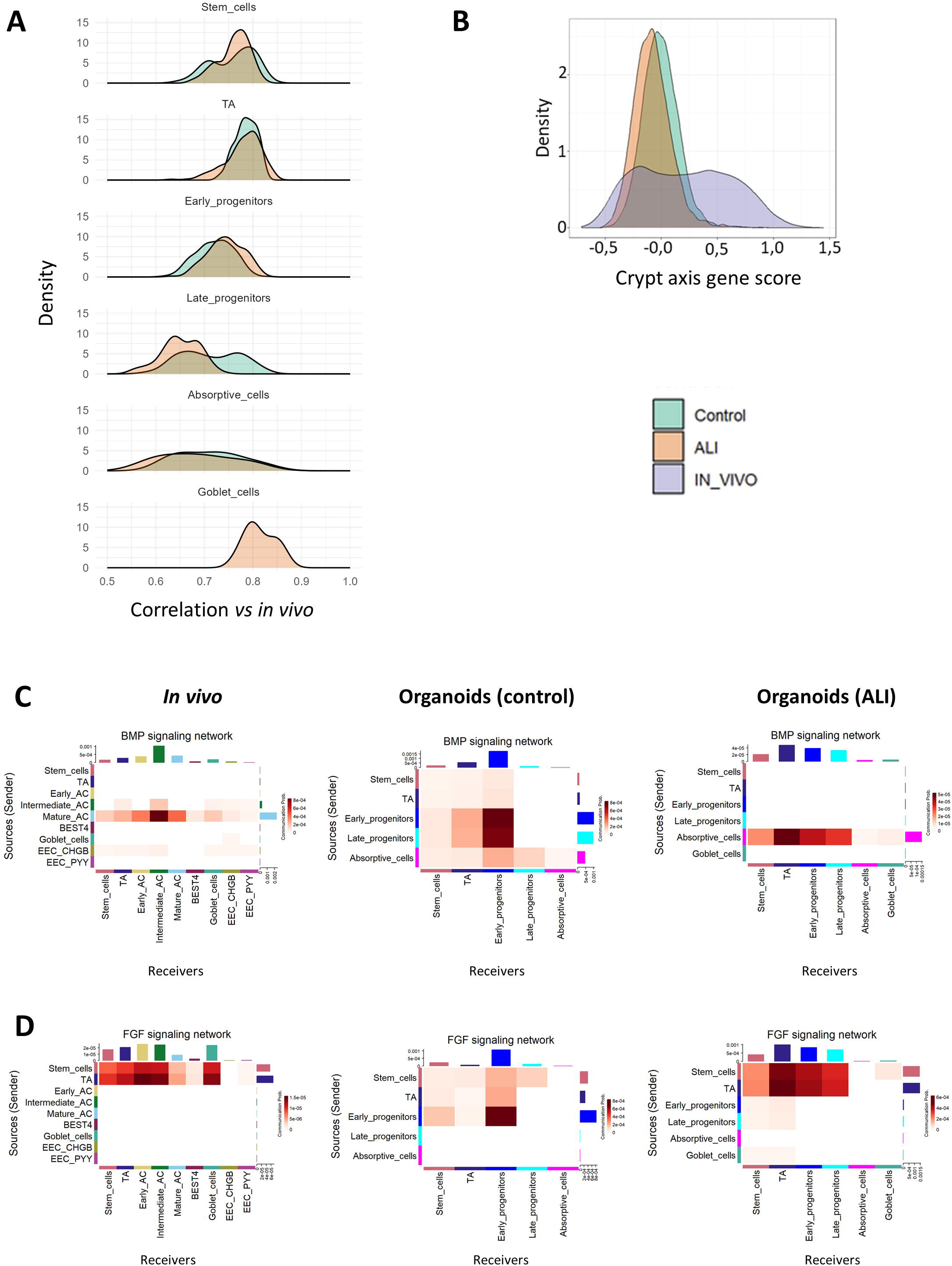
Cellular neighborhood correlation, crypt axis gene scoring, and cell–cell interaction analysis across *in vivo*, control, and ALI conditions. (A) Distribution of the cell neighborhood correlations between *in vivo* and organoid cells in the submerged (control) and the air-liquid interface (ALI) conditions, by cell types. (B) Distribution of crypt axis gene score for the cells of the *in vivo*, control, and ALI conditions. (C) Estimated communication probabilities based on the Bone Morphogenic Protein (BMP) signaling network between each pair of cell types for the *in vivo*, control, and ALI conditions. Senders are presented on the y axis while receivers are presented on the x axis. (D) Estimated communication probabilities based on the Fibroblast Growth Factor (FGF) signaling network between each pair of cell types for the *in vivo*, control, and ALI conditions. AC: absorptive cells, EEC: enteroendocrine cells, TA: transitamplifying cells.

Cell-cell communication analysis was performed to compare predicted cellular interactions in organoid cell monolayers cultured in control or ALI condition with those observed in the caecum epithelium *in vivo*. Tables S7 and S8 provide the estimated communication probability of each pair of cell types for each pathway and each pair of ligand-receptor, respectively. We focused our analysis on bone morphogenic pathway (BMP) and fibroblast growth factor (FGF) signaling pathways, which are critical for cellular communications in the intestinal epithelium (3). Analysis of BMP signaling *in vivo* revealed that mature absorptive cells were the main signal senders of the ligand *BMP2,* while all the other cell types were receivers expressing the receptors *BMPR1A*, *BMPR2*, and *ACVR2A* (Figure 5C and Table S8). A similar cell-cell interaction network for BMP signaling was predicted in organoid cell monolayers cultured in ALI condition. In contrast, all cell types acted both as senders and receivers in the control condition. For FGF signaling *in vivo*, stem cells and TA cells were the primary senders of the ligand *FGF18* towards most cell types through the receptors *FGFR2, FGFR3*, or *FGFR4* (Figure 5D and Table S8). The main senders of FGF signaling were also stem and TA cells in organoid cell monolayers cultured in ALI, while early progenitors were the main senders in the control condition. In contrast with *in vivo* epithelial cells, FGF signals were received by control and ALI and organoid cells through *FGFR2* and *FGFR4* but not *FGFR3*. Overall, our results indicate that culture in ALI increased the resemblance of organoid cells to intestinal epithelial cells *in vivo*.

## DISCUSSION

The development of organoids has opened up new opportunities to study the intestinal epithelium *in vitro* under physiological conditions while reducing the need for animal experimentation (47). However, a close replication of the cellular diversity of the intestinal epithelium is required to ensure the reliability of the results obtained in organoids. In this study, we investigated the cellular heterogeneity of rabbit caecum organoids cultured in a 2D monolayer format that allows access to the apical and basolateral sides of the epithelium (31). To achieve this goal, we used scRNA-seq to compare the transcriptome of each cell type present in organoids with a single-cell transcriptomic atlas of the rabbit caecum epithelium *in vivo* (32).

Organoid cell monolayers are most commonly grown under submerged conditions, *i.e.,* with culture medium on both the apical and basolateral sides (17). In our study, submerged organoid cell monolayers were cultured with an expansion medium on the basolateral side and a growth factor-free medium on the apical side in order to mimic the stromal exposure of epithelial cells to growth factors. Under these culture conditions, the submerged organoid cell monolayers were predominantly composed of stem cells, TA cells and absorptive progenitor cells, which is consistent with previous scRNA-seq analyses of 3D intestinal organoids grown in expansion medium (12, 13, 16, 35). The transcriptome of stem cells and TA cells in rabbit caecum organoid cell monolayers was highly similar to their *in vivo* counterparts, indicating that the stem cell niche was efficiently reproduced *in vitro* in our experimental conditions. Accordingly, most of the organoid cells were predicted to be localized in the lower half of the crypt, where stem cells, TA cells and immature cells are localized *in vivo* (32, 48).

Absorptive cells in rabbit caecum organoid cell monolayers remained mostly in a progenitor state and their transcriptome was dissimilar with *in vivo.* Furthermore, BEST4^+^ cells were not present in organoid cell monolayers although we recently identified that this subset of mature absorptive cells composed up to 5% of the caecum epithelium in rabbits *in vivo* (6, 32). A previous study suggested that complete differentiation of absorptive cells *in vitro* required longer cultivation than other epithelial cell types (49).

Therefore, the 5 days of organoid cell monolayer culture in our study may have been too short to induce full differentiation of absorptive cells. Furthermore, the use of a differentiation medium containing reduced levels of mitogenic factors or the activation of BMP signaling may be required to replicate the crypt-top microenvironment that promotes absorptive cell differentiation *in vivo* (5, 12, 16). For instance, removal of Wnt signaling activators from the culture medium is required for the emergence of BEST4^+^ cells in human intestinal organoids (7, 50). However, the use of a differentiation media may compromise the ability of the organoid cell monolayers to self-renew *in vitro* due to the loss of the stem cell pool. Alternatively, the use of microfluidic “gut-on-a-chip” devices allowing to recreate key features of the epithelial microenvironment (e.g. crypt topography, concentration gradients of growth factor, fluid flows) supported both self-renewal and multilineage differentiation of human organoid cells, including BEST4^+^ cells (49).

Importantly, rabbit caecum organoid cell monolayers lacked cells of the secretory lineage when cultured in submerged conditions. In contrast, we found that the formation of an ALI by removing the apical medium for 2 days was sufficient to induce the differentiation of goblet cells that had a high transcriptional similarity to their *in vivo* counterparts. This increased number of goblet cells in the ALI condition compared to the submerged condition is consistent with several studies performed with human, murine and porcine intestinal organoids (20–24, 28, 51). Additionally, we found that ALI culture led to the emergence of rare EEC, suggesting an overall enhancement of secretory cell differentiation upon apical medium removal. This result is consistent with previous studies showing that ALI conditions promotes the differentiation of EEC in primary human epithelial cell monolayers (21, 24, 52). The increased cellular heterogeneity observed in rabbit caecum organoid cell monolayers in ALI condition was also associated with a closer replication of BMP and FGF signaling cell-cell communications, which both play a pivotal role in epithelial homeostasis (3). Importantly, ALI-induced emergence of secretory cells did not compromise epithelial self-renewal of rabbit caecum organoid cell monolayer since the stem cell pool was maintained and the proportion of TA cells was increased, when compared to immerged conditions. This results is consistent with other studies showing that the switch from immerged to ALI conditions transiently increased proliferation in intestinal epithelial cell monolayers (19). However, the culture of rabbit caecum organoid cell monolayers in ALI conditions did not increase the transcriptomic similarity of absorptive cells when compared to *in vivo*.

The most striking effect of the ALI culture, observed in all cell types, was the strong downregulation of genes involved in hypoxia (53–56). Similar results were obtained in previous studies using ALI culture of intestinal epithelial cell and can be explained by the higher oxygen delivery when cells are directly exposed to air, without liquid at the apical side (29). Since glycolysis is independent of oxygen, the ALI-induced reduction of hypoxia probably explains the lower expression in all cell types of genes coding for enzymes involved in glycolysis (57). Other studies also reported that the culture of intestinal epithelial cells in ALI conditions induces a metabolic switch from glycolysis to mitochondrial oxidative phosphorylation (19, 29, 30, 51). This reduction of hypoxia is directly involved in the pro-differentiation effect of ALI culture since re-submersion of ALI monolayers or reduction of oxygen tension was shown to rapidly reverse epithelial differentiation in organoid cell monolayers (51). Furthermore, the induction of goblet cell differentiation in ALI conditions may be related to the increased oxygen tension that alleviates endoplasmic reticulum stress, which goblet cells are particularly sensitive to, due to their active protein synthesis for mucus secretion (51, 58). This mechanism could also explain the ALI-induced differentiation of EEC, as the endoplasmic reticulum is also involved in hormone secretion (59). The absence of other secretory epithelial cells, namely Paneth cells and Tuft cells, in rabbit organoid cell monolayers was expected, as these cells have not been found by scRNA-seq *in vivo* in the rabbit caecum epithelium (32).

Despite its ability to increase epithelial heterogeneity *in vitro*, the culture of organoid cell monolayers in ALI conditions has some drawbacks. Indeed, under physiological conditions, the apical side of the intestinal epithelium is in a hypoxic state (<1% O_2_), while the basal side is in a normoxic state. On the contrary, the opposite is observed *in vitro* in ALI cultures (23, 60). Furthermore, the absence of fluid on the apical side of epithelial cells complicates treatments aimed at mimicking a luminal exposure to pathogen, metabolites or nutrients. Re-submersion of ALI organoid cell monolayers for apical treatment should be limited to short-term exposure, as this procedure has been shown to reverse epithelial differentiation in organoid cell monolayers as early as 24 h (51).

## CONCLUSION

This study highlights the cellular heterogeneity of rabbit caecum organoid cell monolayers under submerged and ALI conditions. The ALI condition was particularly effective in promoting goblet cell differentiation in a normoxic environment. Furthermore, our results showed that the transition from submerged to ALI conditions induces an energetic metabolic shift in all cell types, likely due to the reduction of hypoxia. Additionally, we showed that the transcriptome of stem cells and TA cells in rabbit organoid cell monolayers was highly similar to their *in vivo* counterpart. In contrast, cells of the absorptive lineage remained largely in a progenitor state. Further improvements in the recapitulation of the intestinal microenvironment are required to allow the phenotype of the intestinal epithelium to be more closely mimicked in cell monolayers derived from organoids.

## Acknowledgments

This work was supported by a grant from the French National Research Agency: ANR-JCJC MetaboWean (ANR-21-CE20-0048). We acknowledge the I2MC cytometry and cell sorting facility (Genotoul-TRI), member of the national infrastructure France-BioImaging supported by the French National Research Agency (ANR-24-INBS-0005 FBI BIOGEN).

## Author contributions

NV and MB designed research;

TM, CM, EL, ER, and MB conducted research;

TM, CC, NV, and MB analyzed data;

TM, NV, and MB wrote the initial draft.

All authors have read and approved the final manuscript.

---

**Figure S1: Quality controls of the organoid single-cell transcriptomics**

(A) Number of expressed genes, counts, and percentage of mitochondrial gene reads per cell and sample before (top panels) and after (bottom panels) filtering for the submerged (control) and air-liquid interface (ALI) conditions.

(B) Uniform Manifold Approximation and Projection (UMAP) of cells from individual samples (inserts) of the control and ALI conditions colored per conditions.

(C) UMAP colored by the number of expressed genes, total counts, or percentage of mitochondrial gene reads per cell for the control and ALI conditions.

(D) UMAP colored by the clusters identified with the *FindClusters* function and expression of the top 10 genes with the highest average log2(fold change) of each cluster. For a given cluster, markers were ordered by decreasing log2(fold change) of the expression between this cluster and the other clusters. Numbers and colored bars on the top indicate cell clusters.

**Figure S2: The air-liquid condition promotes the differentiation of enteroendocrine cells**

Uniform Manifold Approximation and Projection (UMAP) of integrated single cell transcriptomics datasets of caecum epithelial cells *in vivo* and of organoid cell monolayers cultured submerged (control) or with air-liquid interface (ALI), colored by the expression of the enteroendocrine cell markers *CHGA* and *CHGB* with all cells (first row), or only with cells from each condition (other rows). The dotted circle highlights the localization of *in vivo* enteroendocrine cells on the UMAP.

**Table S1. Marker genes of each cell type in the control condition**

List of the marker genes of each cell type (presented in separate tabs) in the organoid cell monolayers under submerged conditions identified by using the *FindAllMarkers* function of Seurat. Columns give the gene name (gene), the *p*-value of the Wilcoxon rank test (pvalue), the adjusted *p*-value (Bonferroni procedure) (p.adjust), the average log2 fold-change (avg_log2FC), the percentage of cells expressing the gene in the cell type indicated in the tab name (pct.1) and the percentage of cells expressing the gene in all other cell types (pct.2). TA: transit amplifying cells.

**Table S2. Marker genes of each cell type in the air-liquid interface condition**

List of the marker genes of each cell type (presented in separate tabs) in the organoid cell monolayers under air-liquid interface conditions identified by using the *FindAllMarkers* function of Seurat. Columns give the gene name (gene), the *p*-value of the Wilcoxon rank test (pvalue), the adjusted *p*-value (Bonferroni procedure) (p.adjust), the average log2 fold-change (avg_log2FC), the percentage of cells expressing the gene in the cell type indicated in the tab name (pct.1) and the percentage of cells expressing the gene in all other cell types (pct.2). TA: transit amplifying cells.

**Table S3. Biological pathways enriched in the marker genes of each cell type in the control condition**

List of biological pathways enriched in the marker genes of each cell type (presented in separate tabs) in the organoid cell monolayers under submerged conditions. Columns give the identifier of the gene ontology (ID), the pathway name (Description), the number of marker genes over the number of genes in the biological pathway (GeneRatio), total number of genes in the biological pathway over the total number of genes expressed in the single-cell RNA-sequencing dataset (BgRatio), the *p*-value (pvalue) and the adjusted *p*-value (p.adjust) (Benjamini-Hochberg procedure) of the Fisher exact test, and the list of marker genes in the biological pathway (geneID), respectively. TA: transit amplifying cells.

**Table S4. Biological pathways enriched in the marker genes of each cell type in the air-liquid interface condition**

List of biological pathways enriched in the marker genes of each cell type (presented in separate tabs) in the organoid cell monolayers under air-liquid interface conditions. Columns give the identifier of the gene ontology (ID), the pathway name (Description), the number of marker genes over the number of genes in the biological pathway (GeneRatio), total number of genes in the biological pathway over the total number of genes expressed in the single-cell RNA-sequencing dataset (BgRatio), the *p*-value (pvalue) and the adjusted *p*-value (p.adjust) (Benjamini-Hochberg procedure) of the fisher exact test, and the list of marker genes in the biological pathway (geneID), respectively. TA: transit amplifying cells.

**Table S5. Differentially expressed genes between groups in each cell type**

List of the differentially expressed genes (DEGs) between conditions (submerged (control) vs air-liquid interface (ALI)) in each cell type (presented in separate tabs) identified by fitting Negative Binomial generalized linear models on pseudo-bulk data in each cell type independently. The columns give the gene name, the log2(fold change) (logFC), the log counts per million (CPM) reads (logCPM), the likelihood ratio (LR), the *p*-value (LR test) (pvalue), the adjusted *p*-value (Benjamini-Hochberg procedure) (p.adjust), and the over or under-expressed state of the gene in the “ALI” condition versus the “control” condition (state), respectively. TA: transit amplifying cells.

**Table S6. Biological pathways enriched in differentially expressed genes between conditions in each cell type.**

List of biological pathways enriched in differentially expressed genes (DEGs) between conditions (submerged (control) vs air-liquid interface (ALI)) per cell type. Columns give the identifier of the gene ontology (ID), the pathway name (Description), the number of marker genes over the number of genes in the biological pathway (GeneRatio), total number of genes in the biological pathway over the total number of genes expressed in the single-cell RNA-sequencing dataset (BgRatio), the *p*-value (pvalue) and the adjusted *p*-value (p.adjust) (Benjamini-Hochberg procedure) of the fisher exact test, and the list of DEGs in the biological pathway (geneID). TA: transit amplifying cells.

**Table S7. Cell-cell communication probabilities across cellular pathways for the *in vivo*, the control and the air-liquid interface conditions**

List of the communication probabilities between all pairs of cell types for multiple signaling pathways for the submerged (control) and air-liquid interface conditions (presented in separate tabs). Columns provide the names of the senders (Senders), receivers (Receivers), the pathways in which the senders and the receivers are involved in (Pathway), and the estimated probability of communication between the senders and the receivers (Probability of communication).

**Table S8. Cell-cell communication probabilities based on the expression of ligand-receptor pairs for the *in vivo*, the control and the air-liquid interface conditions**

List of the communication probabilities between all pairs of cell types based on several pairs of ligand-receptors for the submerged (control) and air-liquid interface (presented in separate tabs). Columns provide the names of the senders (Senders), the receivers (Receivers), the pathways in which the senders and the receivers are involved in (Pathway), the ligand-receptor pairs (Ligand-receptor pairs) and the estimated probability of communication between the senders and the receivers (Probability of communication).

