## Supplementary figures and images for "Single cell transcriptomics reveals that air-liquid interface culture promotes goblet cell differentiation and inhibits glycolysis in cell monolayers derived from rabbit caecum organoids"

### Figure S1

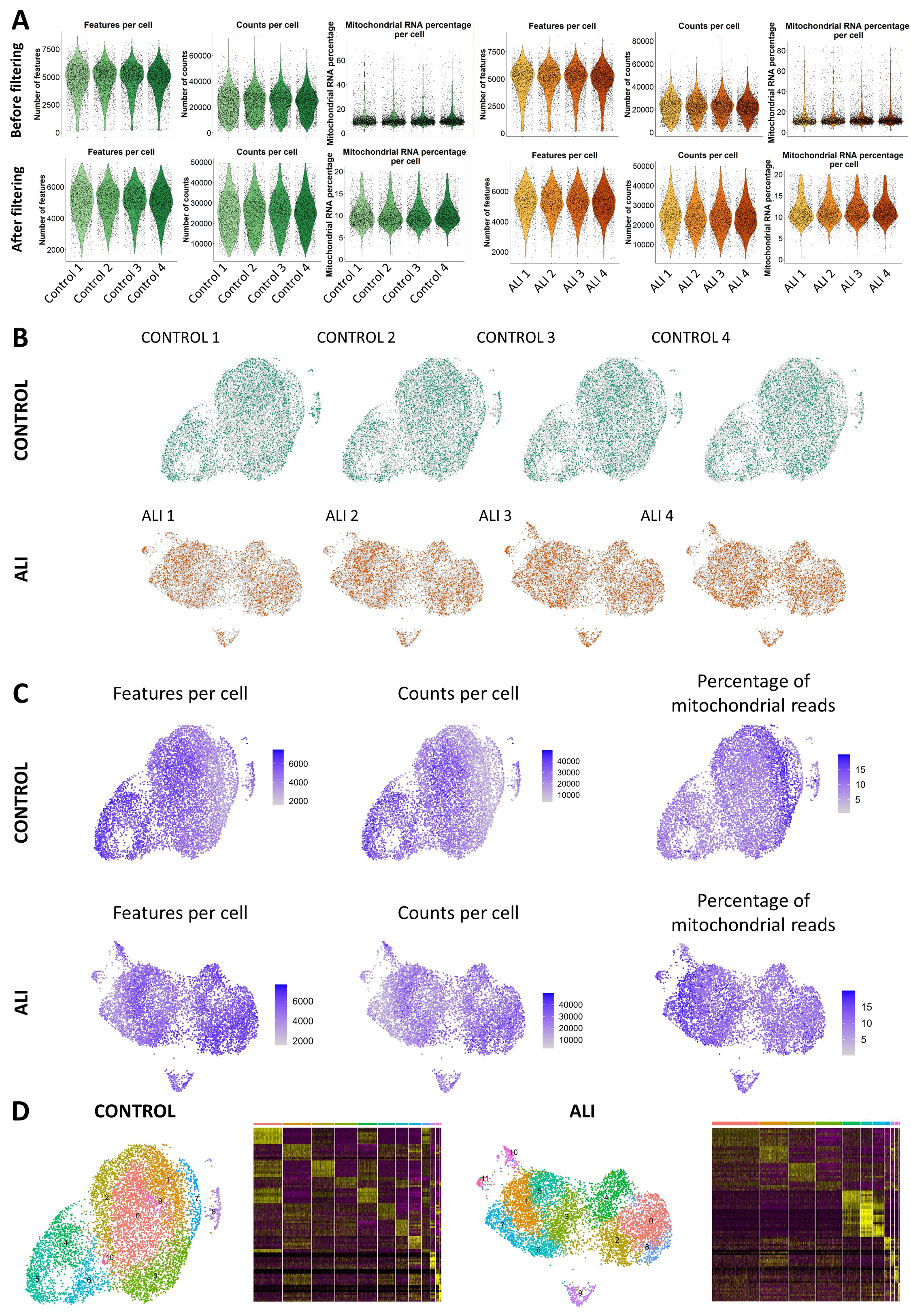

### Figure S2

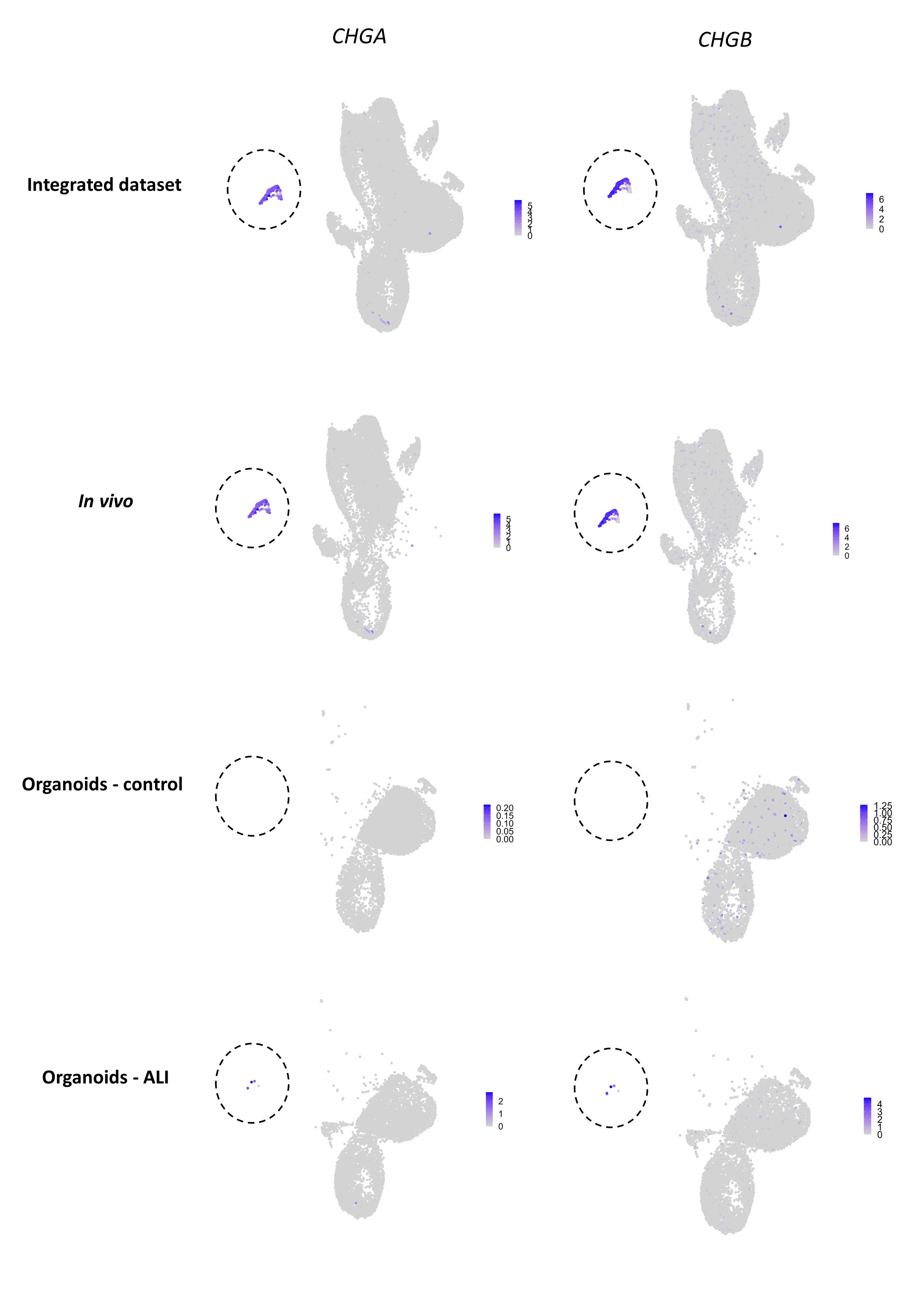
